# Branched Oncolytic Peptides Target HSPGs, Inhibit Metastasis, and Trigger the Release of Molecular Determinants of Immunogenic Cell Death in Pancreatic Cancer

**DOI:** 10.1101/2024.09.03.610940

**Authors:** Alessandro Rencinai, Eva Tollapi, Giulia Marianantoni, Jlenia Brunetti, Tania Henriquez, Alessandro Pini, Luisa Bracci, Chiara Falciani

**Affiliations:** Department of Medical Biotechnology, University of Siena, Siena, Italy

**Keywords:** Immunogenic cell death, pancreatic cancer, oncolytic peptides

## Abstract

Immunogenic cell death (ICD) can be exploited to treat non-immunoreactive tumors that do not respond to current standard and innovative therapies. Not all chemotherapeutics trigger ICD, among those that do exert this effect, there are anthracyclines, irinotecan, some platinum derivatives and oncolytic peptides. We studied two new branched oncolytic peptides, BOP7 and BOP9 that proved to elicit the release of damage-associated molecular patterns DAMPS, mediators of ICD, in pancreatic cancer cells. The two BOPs selectively bound and killed tumor cells, particularly PANC-1 and Mia PaCa-2, but not cells of non-tumor origin such as RAW 264.7, CHO-K1 and pgsA-745. The cancer selectivity of the two BOPs may be attributed to their repeated cationic sequences, which enable multivalent binding to heparan sulfate glycosaminoglycans (HSPGs), bearing multiple anionic sulfation patterns on cancer cells. This interaction of BOPs with HSPGs not only fosters an anti-metastatic effect *in vitro*, as demonstrated by reduced adhesion and migration of PANC-1 cancer cells, but also shows promising tumor-specific cytotoxicity and low hemolytic activity. Remarkably, the cytotoxicity induced by BOPs triggers the release of DAMPs, particularly HMGB1, IFN-β and ATP, by dying cells, persisting longer than the cytotoxicity of conventional chemotherapeutic agents such as irinotecan and daunorubicin. An *in vivo* assay in nude mice showed an encouraging 20% inhibition of tumor grafting and growth in a pancreatic cancer model by BOP9.

## INTRODUCTION

The primary explanation for the poor prognosis often associated with pancreatic cancer, particularly pancreatic ductal adenocarcinoma (PDAC), is its immunologically inert nature. PDAC is characterized by a prevalence of immunosuppressive infiltrating cells (1-3) and it also has a low mutational burden, resulting in low levels of neoantigens and a compromised capacity for T cell recognition of tumors (4-5). Consequently, immune checkpoint inhibitors (ICIs), a cornerstone of immunotherapy, have shown little efficacy in PDAC patients (6). Successful stimulation of the tumor immune microenvironment is required for an effective PDAC antitumor immune response (7).

Immunogenic cell death (ICD) is a regulated type of cell death that drives antigen-specific immune responses culminating in immunological memory (8). Various endogenous adjuvant signals, referred to as damage-associated molecular patterns (DAMPs), are released by malignant cells during immunogenic stress or cell death, stimulating the maturation of antigen-presenting cells (9). DAMPs include endoplasmic reticulum proteins, calreticulin (CRT), ATP, high mobility group box 1 (HMGB1), and type I interferon (type I IFN). These DAMPs serve collectively as “eat me” and “find me” signals, attracting antigen-presenting cells to sites of immunogenic cell death induction. This process stimulates the uptake, processing, and cross-presentation of tumor-associated antigens, ultimately eliciting an adaptive immune response (10).

ICD-inducing anticancer agents are extremely interesting, especially for “cold cancers”, such as pancreatic cancer, because they exhibit “dual-action”: while directly killing most tumor cells, dying cancer cells act as a sort of vaccine that triggers a specific immune response aimed at eradicating the remaining cancer cells.

Indeed, only some chemotherapeutics can induce intracellular stress pathways that activate a DAMP release in dying cancer cells. Conventional chemotherapeutics that do so include cyclophosphamide, anthracyclines and some platinum derivatives (10-11). Photodynamic therapy (PDT) (12) and radiotherapy with γ irradiation (13) have also proved effective in triggering ICD.

The literature contains a vast number of examples of peptides with cytolytic sequences and selectivity for cancer cells (14-16). Most are derived from natural host defence peptides, highly conserved peptides, synthesized by almost all living organisms. They generally consist of 10-30 amino acids that confer a global amphipathic conformation with a positive net charge originating from the high prevalence of cationic (e.g. Lys, Arg) and hydrophobic (e.g. Ala, Val, Gly) amino acids, that allow interaction with biological membranes. These oncolytic peptides can also elicit immunogenic cell death by triggering release of ICD-related DAMPs (17-19). Many oncolytic peptides are reported to interact with cell surface heparan sulfates of proteoglycans and to penetrate the plasma membrane (20-22). These interactions occur between the cationic, basic amino acids on the peptides and the negatively charged sulfates or carboxyl groups of the glycosaminoglycan (GAG) portion of heparan sulfate glycosaminoglycans (HSPGs) and/or sialic acid, typically increased in cancer cells (17). HSPGs can therefore function as the initial selective anchoring site for these oncolytic peptides.

One of the limitations of using short linear peptides in clinical settings is their low half-life in vivo due to rapid degradation by peptidases and proteases. This restricts their utility to intratumoral administration (23). Despite the very promising properties of oncolytic peptides being well-established, only one clinical trial has been published (24) where patients with metastatic soft tissue sarcoma were treated with the combination of an intratumoral injection of oncolytic peptide and adoptive T-cell therapy, in a phase one study (24).

Tetrabranched dendrimeric peptides have much better resistance to hydrolysis than their monomeric counterparts. This feature is due to their steric hindrance that limits interaction with the cleavage site of peptidases (25-27). Besides, branched peptides enable greater binding avidity because they form polyvalent bonds. The branched structure, with multiple active sequences on the same molecule, also determines higher local concentrations than can’t be achieved with linear homologs (26-27).

We accordingly synthesized and assessed the cytotoxic specificity of two oncolytic tetrabranched peptides against cancer cells. We also investigated their potential antimetastatic effects through interaction with HSPGs, and their ability to induce immunogenic cell death. The efficacy of these peptides was also evaluated in a murine model of pancreatic cancer.

## MATERIALS AND METHODS

### Peptide synthesis

Linear peptides L7 and L9 and branched oncolytic peptides BOP7 and BOP9 were synthesized on solid-phase by standard Fmoc chemistry on a Syro multiple-peptide synthesizer (MultiSynTech, Witten, Germany). The branched peptides were synthesized on a TentaGel S RAM resin (Rapp Polymere, Tübingen, Germany). For the branched peptides the first two coupling steps were carried out with Fmoc-Lys(Fmoc)-OH to create the branching core. In the synthesis of biotin-conjugate peptides, Fmoc-Lys(Biotin)-OH was used in the initial coupling step, followed by Fmoc-PEG4-OH in the subsequent step, both preceding introduction of the branching core. Sidechain-protecting groups were 2,2,4,6,7-pentamethyldihydrobenzofuran-5-sulfonyl for R, t-butoxycarbonyl for K, Trityl for Q and t-butyl for S. The final product was cleaved from the solid support, deprotected by treatment with TFA containing triisopropylsilane and water (95/2.5/2.5) and precipitated with diethyl ether. Crude peptide was purified by reversed-phase chromatography on a Phenomenex Jupiter C18 column (300 Å, 10 mm, 250, 610 mm), using 0.1% TFA/water as eluent A and methanol as eluent B, in a linear gradient from 80% A to 50% A in 30 min.

Final peptide purity and identity were confirmed by reversed-phase chromatography on a Phenomenex Jupiter C18 analytical column (300 Å, 5 mm), L7=16.28 min; L9=14.24 min; BOP7 =20.04 min; BOP9=16.44 min, BOP7-Bio=21.33 min, BOP9-Bio=19.43 min and by mass spectrometry with a Bruker Daltonics Ultraflex MALDI TOF/TOF: L7 M+(found) = 1274.159; L9 M+(found) = 1289.085; BOP7 M+(found) = 5.425.857; BOP9 M+(found) = 5.486.092; BOP7-Bio M+(found) = 6.028.439; BOP9 M+(found) = 6.088.041.

### Stability to serum proteolysis

A pool of sera from healthy volunteers (n=4) was diluted 25% using RPMI 1640 medium. Each peptide was incubated at 37°C for 4 and 16 hours at different concentrations in the diluted serum. Trichloroacetic acid (TCA), diluted 15% in water, was then added to the samples which were then centrifuged at 12,000 × g. The resulting supernatant was diluted 20% with 0.1% TFA in water and used for HPLC analysis. The HPLC spiked peaks were collected and used for MALDI-TOF analysis. Time-zero HPLC and MS-spectroscopy spectra were obtained immediately after mixing each peptide with 25% serum. The presence of the intact peptide was confirmed by MALDI mass spectrometry.

### Cell cultures

Cell lines, purchased from The Global Bioresource Center (ATCC, Rockville, MD, USA), were maintained at 37°C in a 5% CO_2_ atmosphere.

PANC-1 and Mia PaCa-2 human pancreas adenocarcinoma, RAW 264.7 murine macrophages, CHO-K1, chinese hamster ovary cells and pgsA-745, chinese hamster ovary cell mutant deficient in xylosyltransferase (UDP-D-xylose: serine-1,3-D-xylosyltransferase), were grown in their recommended medium: Dulbecco’s Modified Eagle’s Medium (DMEM). Medium was supplemented with 10% fetal bovine serum (FBS), 200 μg/mL glutamine, 100 μg/mL streptomycin and 60 μg/mL penicillin. The culture medium of Mia PaCa-2 was further supplemented with 2,5% horse serum (HS).

PANC-1-*luc2* cells, luciferase-expressing cell line isolated were obtained from PANC-1 cells, transfected with pGL4.51[luc2/CMV/Neo] vector (Promega, Madison, WI, USA) using Lipofectamine™ 3000 Reagent (Thermofisher), following the manufacturer’s instruction.

### Flow cytometry

PANC-1, Mia PaCa-2, RAW 264.7 and pgsA-745 cells were seeded 2 × 10^5^ cells/well in 96-well U-bottom plates. They were incubated with different concentrations (10 μM, 2 μM, and 0.4 μM) of biotinylated BOP7 and BOP9 for 30 min at room temperature, followed by incubation with Streptavidin-FITC diluted 1:1000 (Sigma Aldrich, St. Louis, MO, USA). All dilutions were performed in PBS, containing 5 mM EDTA and 1% BSA. For BOP binding, 7000 events were evaluated in a Guava easyCyte Flow Cytometer (Millipore). For the competition experiment with heparin, BOP7 and BOP9 were used at 10 μM in the presence of heparin at 5 and 20 μM. Experiments were repeated at least twice and each group evaluation was conducted in duplicate. The results were analysed by FCS Express 6 Flow cytometry software.

### BOP binding to heparin -ELISA

BOP7 and BOP9 were diluted in carbonate buffer (pH 9) and used at concentrations of 10 μg/mL, 5 μg/mL and 1 μg/mL to coat a 96-well ELISA strip plate in which uncoated wells were used as negative controls. The plate was then sealed and incubated overnight at 4°C, then saturated with milk 3% in PBS. The plate was incubated with heparin-biotin sodium salt (Sigma Aldrich) diluted in PBS-BSA 0.3% to 5 μg/mL. After 30 minutes incubation and washing, streptavidin-POD (Sigma Aldrich) (1:500 in PBS-milk 0.3%) 100 μL/well was added and incubated in the dark for 30 minutes at 30°C. After washing, 150 μL/well of substrate solution (phosphocitrate buffer, TMB, DMSO, glycerol and H_2_O_2_) was added and incubated for 5 minutes. The reaction was quenched with 50 μL/well of HCl 1M and the plate was read at 450 nm and 650 nm using a microplate spectrophotometer (Multiskan, Thermo Scientific, Waltham, MA, USA). The data was analysed using GraphPad Prism version 9.5.0 software.

### Cell viability assays

PANC-1, PANC-1-luc2, Mia PaCa-2 or CHO-K1 cells were seeded 5 × 10^4^ cells/well in a standard 96-well plate and incubated overnight at 37°C, 5% CO_2_ atmosphere. Each well was treated with 200 μL BOP7 or BOP9 at different concentrations (from 100 μM to 0.08 μM) and incubated for 24 hours. PANC-1 and Mia PacCa-2 were also tested with linear peptides L7 and L9 at concentrations ranging from 781 μM to 0.5 μM under the same conditions. Growth inhibition was assessed with a MMT assay (3-(4,5-dimethylthiazol-2-yl)-2,5-diphenyltetrazolium bromide) (Sigma Aldrich). Optical density was measured at 595 nm and 650 nm with a microplate spectrophotometer (Multiskan, Thermo Scientific). Experiments were repeated twice in quintuplicate. Cell viability was measured by comparing values of treated and untreated cells. IC50 values were calculated by non-linear regression analysis using GraphPad Prism version 9.5.0 software. Untreated cells showed 100% cell viability.

### Hemolysis test

Whole human blood was centrifuged at 3500 rpm for 10 minutes. After several washings, isolated cells were diluted 1:50 in PBS and added to a flat 96-well plate (100 μl/well). Serial dilutions of the peptides, from 160 μM to the lowest at 0.25 μM, were added. 100% hemolysis was obtained with 1% TritonX 100 in PBS. The plate was incubated at 37°C for 30 minutes, centrifuged at 2000 rpm for 5 minutes and the supernatant analysed using a plate spectrophotometer (Multiskan, Thermo Scientific) at 405 nm and 490 nm wavelengths.

### Adhesion assay

PANC-1 cells were seeded at a concentration of 2 × 10^6^ cell/ml in a flat 96-well plate. Immediately after, 10, 1 and 0,1 μM BOP7 and BOP9 were added to the plate. Untreated cells were used as control. Cells were incubated at 37°C for 4h and then fixed with 4% PFA-PBS (Sigma Aldrich). The plates were then washed and stained with 100 μl/well 0.1% Crystal Violet (Sigma Aldrich). After treatment with 10% acetic acid the signal was read at 595 nm (Multiskan, Thermo Scientific).

### 2D Migration assay

Two-well silicone inserts (Ibidi GmbH Gräfelfing Germany) were used for this assay. 3 × 10^4^ PANC-1 cells/well were plated in a final volume of 70 μl of culture media for each side of the insert. After 24h incubation at 37°C, the inserts were removed and BOP7 and BOP9, at the concentration of 1μM and 10μM, were added. The closure of the gap was monitored with a DFC 7000T microscope (Leica) taking pictures every 30 minutes for 16h. The remaining gap was measured with ImageJ.

### 3D Migration assay

The 24-well transwell inserts (Sarstedt Nümbrecht, Germany) were handled according to the manufacturer’s instructions. The inserts were coated with Collagen type I (167 μg/ml) (Corning, NY USA) overnight at 37°C. PANC-1 cells were seeded in the upper chamber at a concentration of 5× 10^5^ cells/ml together with BOP7 and BOP9 at 0.1 μM, 1 μM and 10 μM dissolved in DMEM. 600 μl complete media was placed in the lower chamber, and cells were incubated for 24h at 37°C. Cells on the upper chamber membrane were then swabbed and the insert was fixed with 4% PFA-PBS and stained with 0.1% Crystal Violet (Sigma Aldrich). Images were taken in bright field mode with a Leica TCS SP5 microscope. The graph was obtained by measuring the color of each well with ImageJ and normalizing all readouts to the untreated wells.

### HMGB1 release

PANC-1 cells were seeded 5 × 10^4^ cells/well in a standard 96-well plate and incubated overnight at 37°C in a 5% CO_2_ atmosphere. Then they were treated with BOP7, BOP9, daunorubicin hydrochloride (Sigma Aldrich) and irinotecan hydrochloride (Sigma Aldrich) at different concentrations (from 300 μM to 2 μM in 200 μL) and left for 24 hours at 37°C. The following day, release of HMGB1 in the culture medium was assessed with an HMGB1 ELISA kit (IBL International, Hamburg, Germany) in the PANC-1 supernatants, according to the manufacturer’s instructions. Optical density was measured at 450 nm with a microplate spectrophotometer (Multiskan, Thermo Scientific). Data was analysed using GraphPad Prism version 9.5.0 software.

### IFN-β release ELISA

PANC-1 cells were seeded 5 × 10^4^ cells/well in a 96-well plate and incubated overnight at 37°C in a 5% CO_2_ atmosphere. Cells were then treated with BOP7, BOP9, daunorubicin hydrochloride (Sigma Aldrich) and irinotecan hydrochloride (Sigma Aldrich) at different concentrations (from 300 μM to 2 μM) and left for 24 hours at 37°C.

At the same time, a DuoSet ELISA plate (R&D Systems Inc, Minneapolis, MN, USA) for the detection of IFN-β was prepared following the manufacturer’s instructions. The plate was sensitized with 100 μL/well of capture antibody for 24 hours. PANC-1 supernatants were collected and centrifuged for 5 minutes at 1200 rpm, added to the DuoSet plate at 100 μL/well and incubated for 2 hours at room temperature. The detection antibody conjugated with streptavidin-peroxidase was then added. After adding the substrate solution, the plate was read at 450 nm using a microplate spectrophotometer (Multiskan, Thermo Scientific). Data was analysed using GraphPad Prism version software.

### Extracellular ATP bioluminescence assay

PANC-1 cells were seeded (10 × 10^3^ cells/well) in an opaque-walled 96-well plate and incubated overnight at 37°C in a 5% CO_2_ atmosphere. The following day, each well was treated with 150 μL BOP7, BOP9, daunorubicin hydrochloride (Sigma Aldrich) or irinotecan hydrochloride (cod. PHR2717, Sigma Aldrich) at different concentrations (from 50 μM to 2 μM). At the same time, each well was incubated with 50 μL RealTime-Glo Extracellular ATP Reagent Substrate (Promega). The plate was incubated at 37° and bioluminescence was measured at 560 nm at different time intervals with a VICTOR NIVO 5S luminometer (Perkin Elmer, Waltham, MA, USA). Data was analysed using GraphPad Prism version 9.5.0 software.

### Mouse model of metastases

Female BALB-c nu/nu 5–6 weeks old mice were obtained from Charles River Laboratories Italia s.r.l. All mice were housed in cages in a pathogen-free animal facility according to local and European Ethical Committee guidelines. Tumor cells PANC1-*luc2* were harvested, washed and injected into the caudal vein (iv) (0.5-1 × 10^6^ PANC1-*luc2* cells per mouse/100 μL PBS) and the animals were randomized into two groups of five: 1) Untreated; 2) Treated with BOP9, intraperitoneal 20 mg/kg in 150 μL saline solution.

Treatments were administered every day for the first 5 days, starting 4 hours after the transplant, followed by 2 days of rest, and then 3 treatments again for a total of 18 treatments over 32 days. Tumor growth was measured using an IVIS Spectrum Imaging System (IVIS, PerkinElmer). Mice were anaesthetized by vaporized isoflurane and injected subcutaneously (s.c.) with D-luciferin (2.5 mg/mouse Promega) in 100 μl PBS vehicle. Images were taken 15 min after Lucifer injection and photon emissions were collected. Bioluminescence was calculated in regions of interest (ROIs) and intensity was recorded for each tumor. Images of the mice were taken every 2-4 days. Treatment and imaging schedule are reported in Figure 5A. The mice were monitored for 32 days then euthanized. Authorization by the Local EC n. 062018.

### Statistical analysis

The results were analysed by Student’s t-test using GraphPad Prism (GraphPad, San Diego, CA, USA). When comparing more than two groups, one-way analysis of variance (ANOVA) followed by the Dunnet multiple comparisons test was used to detect significant differences between treatment group means. The data is reported with standard deviation (SD).

## RESULTS

### Synthesis of branched peptides

The peptides were synthesized as tetrabranched structures having four oligomers linked to a branching three-lysine core (Figure 1). The sequences were chosen from a comprehensive open-access database containing information on amino acid sequences, chemical modifications, 3D structures, bioactivities and toxicities of peptides (DBAASP v3.0). The peptides of the database derive from host defense peptides or non-natural libraries of peptides. We selected two sequences LLKKKFKKLQ (L7) and KKKLKFKKLQ (L-9) because they had already demonstrated activity against breast carcinoma, cervical hepatocellular and pancreatic cancer (19). Both sequences were synthesized in linear (L-7 and L-9) and branched form (BOP7 and BOP9) (Figure 1).

**Figure 1.**
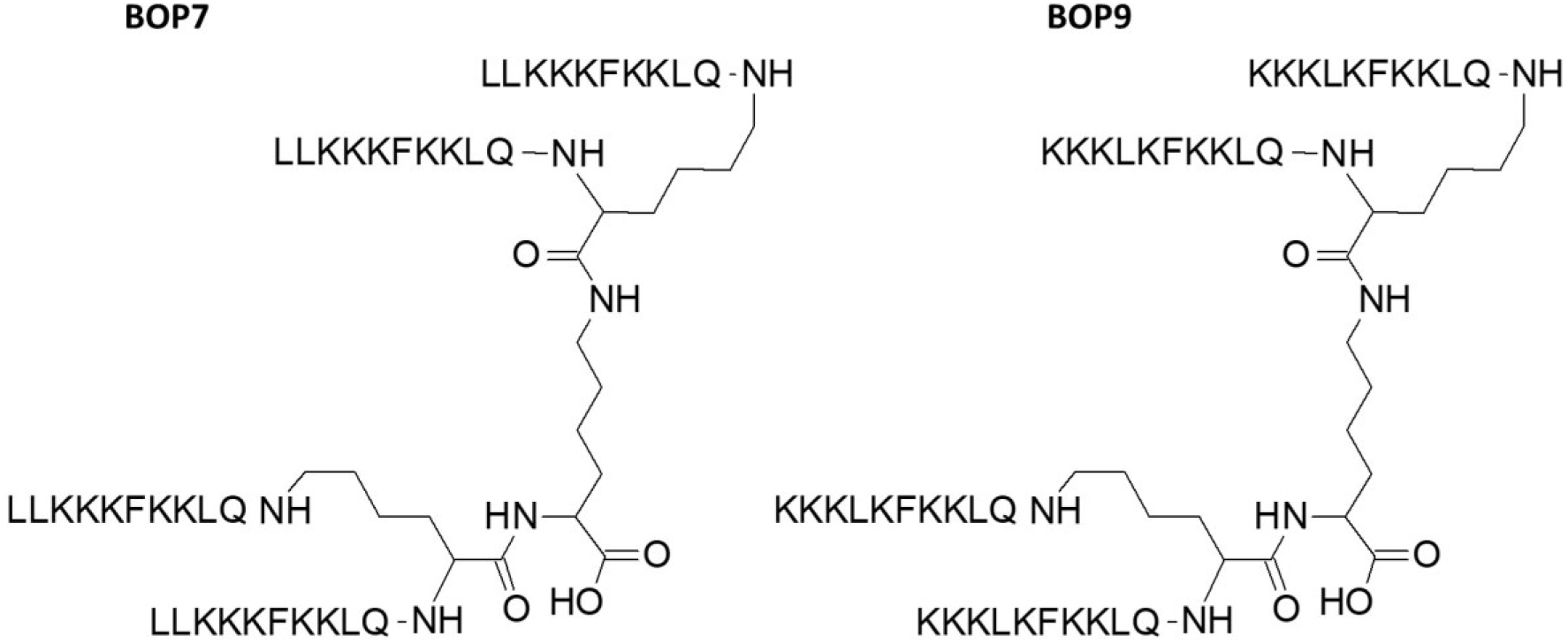
Structure of branched oncolytic peptides (BOPs). The three-lysine core allows assembly of four identical copies of the same sequence

### Stability of BOPs to blood proteases

Serum was taken as a model of a complex mixture of different proteases to test the stability of BOPs to hydrolysis and compare it with the linear analogues L7 and L9. Tetrabranched peptides were incubated in serum for 4 hours and 16 hours at 37°C. HPLC analysis followed by mass spectrometry identified a peak indicating intact peptide (Figure 2) up to 16 h. BOP7 and BOP9 were still detectable after 16 h of incubation in serum, differently from the linear analogues that were hydrolyzed before the 16 h. The very short half-life of the linear analogues prevents their further development.

**Figure 2.**
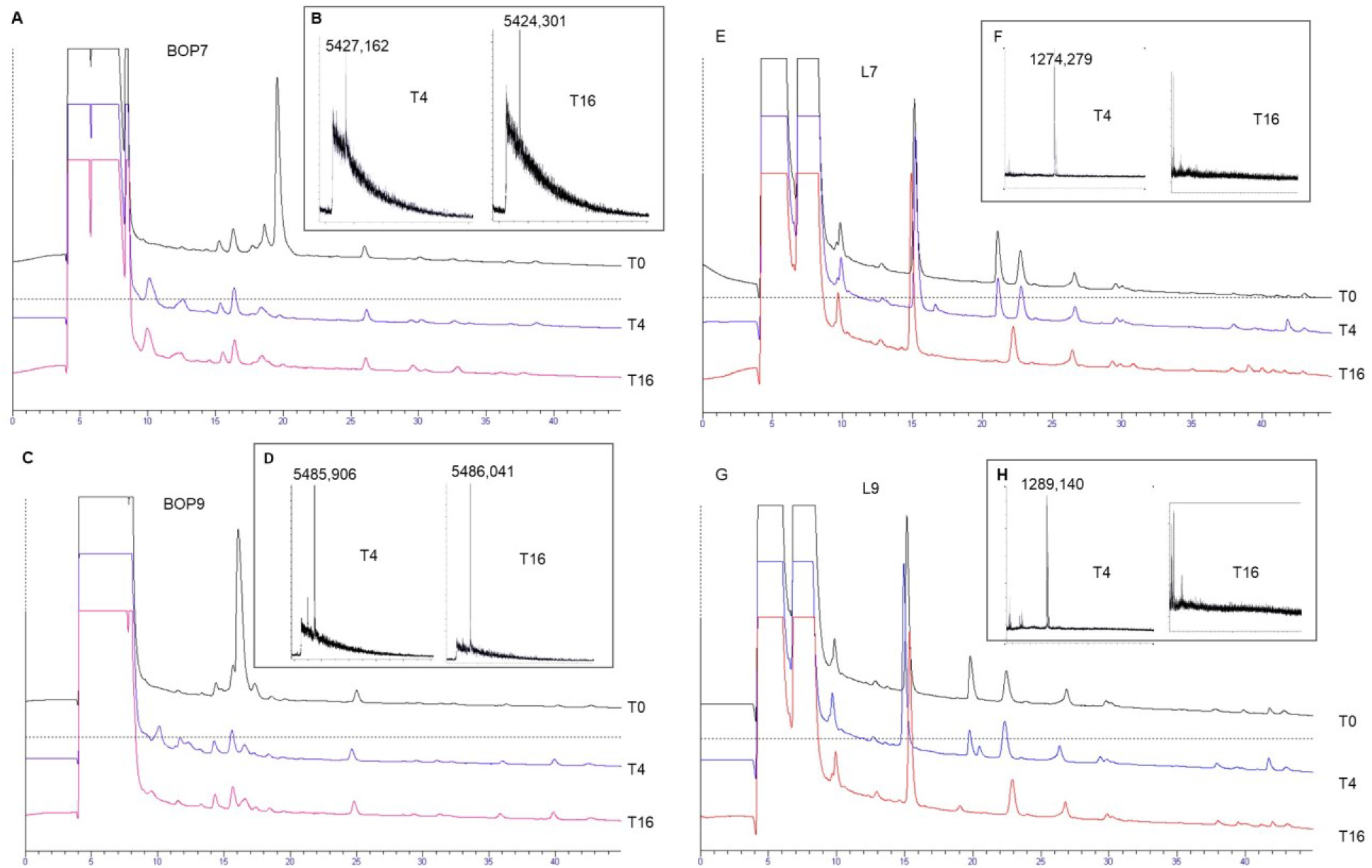
Stability of the tetrabranched peptides to proteolysis. A, C, E, G) HPLC profiles of the serum with spikes of the peptides, BOP7, BOP9, L7 and L9 monitored after 4h and 16h of incubation; B, D, F, H) MALDI-TOF mass spectrometry confirming the presence or absence of the intact peptides.

### Specific binding of BOPs to cancer cell lines through HSPGs

PANC-1 and Mia PaCa-2 pancreatic cell lines were chosen to test the ability of BOPs to bind human cancer cell lines. FACS analysis confirmed that BOPs bind both cell lines in a dose-dependent manner (Figure 3A-B and E), with stronger binding at 2 μM and 10 μM for both BOP7 and BOP9.

**Figure 3.**
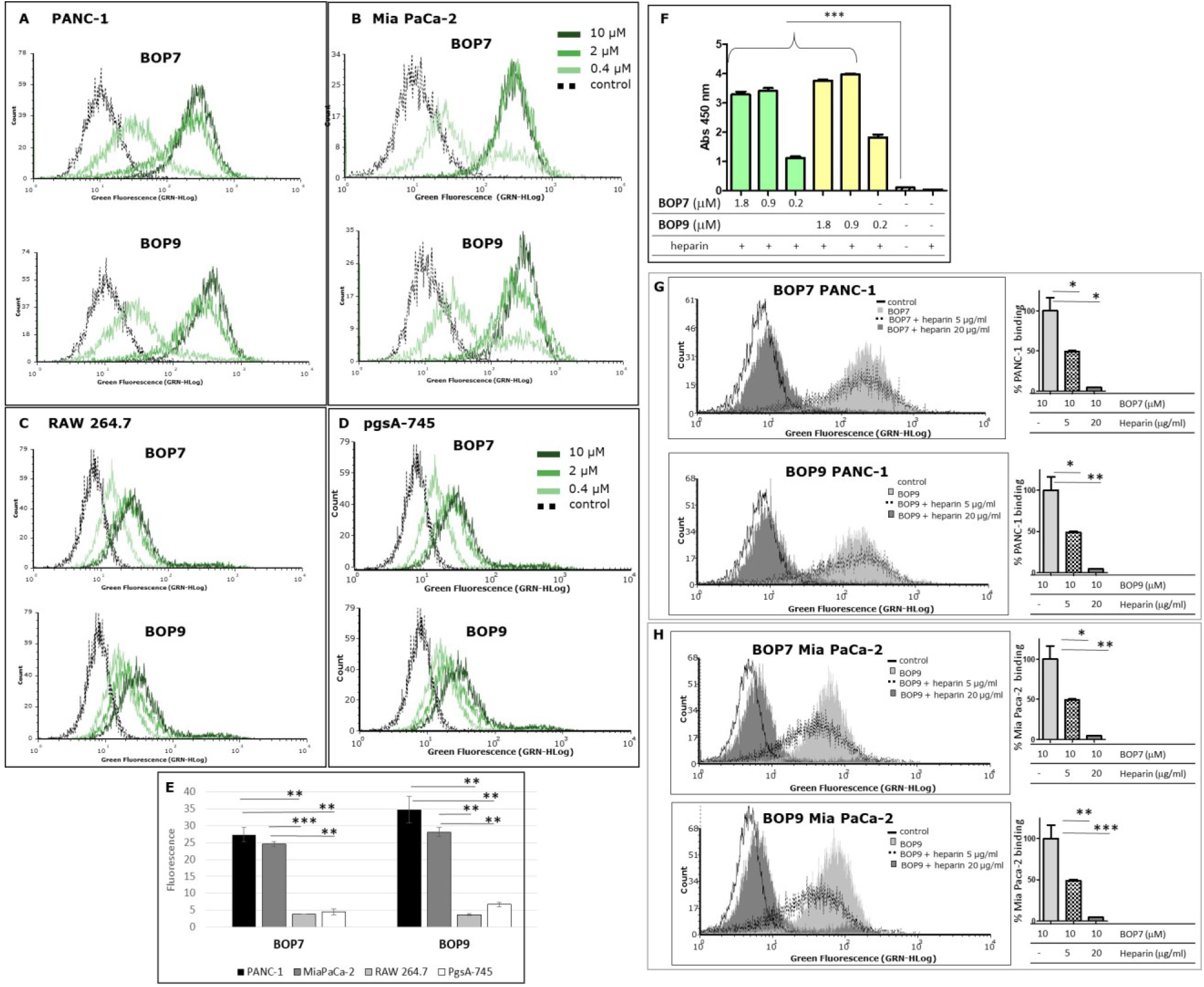
**A-B**) Flow cytometry analysis of: pancreatic cancer cells, PANC-1 and Mia PaCa-2; **C-D**) cells of non-cancer origin, RAW 264.7, murine macrophages and pgsA-745 chinese hamster ovary cell mutant deficient in xylosyltransferase. **E)** Quantification of fluorescence intensity of cells treated with 10 μM BOPs, each repeated value (n=2) was averaged and divided by the value of the untreated cells; **F**) ELISA of binding of BOPs to biotinylated heparin; *** p < 0.0001 (n = 6), one-way ANOVA and Dunnet post-test. **G**) and **H**) Flow cytometry analysis of BOP7 and BOP9 binding to PANC-1 and Mia PaCa-2 cells in the presence of 20 or 5 μg/ml heparin. On the right side of each graph, the quantification of fluorescence intensity, as % of BOPs binding (n=2).

To assess binding specificity, immortalized mammalian cell lines of non-cancer origin were analysed under the same flow-cytometry conditions. RAW 264.7 macrophage cells showed low BOP7 and BOP9 binding, exhibiting fluorescence intensities more shifted to the isotypic control, compared to PANC-1 and Mia PaCa-2 (Figure 3C and E).

Since cationic peptides preferentially bind negatively-charged targets, among which the sulfated GAG chains of HSPGs could be preferential, we chose the pgsA-745 cell line to control selectivity. These cells are mutants of chinese hamster ovary cells deficient in xylosyltransferase, the enzyme responsible for the first sugar coupling in the synthesis of the polysaccharide chains of HSPGs, therefore they completely lack the GAG chains (28). Interestingly, BOP7 and BOP9 showed poor binding activity on pgsA-745 (Figure 3D and E), suggesting that they may target the heparan sulfate chains of HSPGs.

Given the cationic nature of BOPs and their poor binding to pgsA-745, an ELISA assay was conducted to assess their capacity to bind heparin, a commonly used model for studying HSPGs due to their similar chemical structure.

The results demonstrated that both BOPs showed statistically significant binding to heparin across the range of concentrations tested (1.8, 0.90, and 0.2 μM), compared to the control groups, which included uncoated wells and wells lacking heparin. The binding capacity of BOP7 and BOP9 to heparin reached a maximum and plateaued at 0.90 μM (5 μg/mL) (Figure 3F).

To evaluate the ability of BOP7 and BOP9 to bind to HSPGs on the plasma membrane of PANC-1 and Mia PaCa-2 cells, flow cytometry analysis (FACS) was performed employing heparin as an antagonist at two concentrations (5 μg/ml and 20 μg/ml) during the cell binding assay. At the higher concentration (20 μg/ml), heparin completely antagonized the binding of BOPs to PANC-1 and Mia PaCa-2 cells (Figure 3G-H), reinforcing the hypothesis that BOPs bind HSPGs on the membranes of cancer cells.

### Cytotoxicity of BOPs for pancreatic cancer cells

The specific cytotoxicity of BOPs against cancer cells was tested in PANC-1 and Mia PaCa-2 cell lines and IC50 were measured in a 24 h treatment experiment. The two BOPs showed micromolar IC50 values. Specifically, BOP7 and BOP9 showed an IC50 of 1.67 × 10^−6^ M and 8.3 × 10^−6^ M against PANC-1, respectively; and against Mia PaCa-2 the IC50 was 2.73 × 10^−5^ M for BOP7 and and 1.78 × 10^−5^ M for BOP9 (Figure 4A and B). Interestingly, the two linear analogues showed no cytotoxic activity against PANC-1 and Mia PaCa-2 (Figure 4C and D, respectively) in 24h, confirming that the branched design increases biological activity.

**Figure 4.**
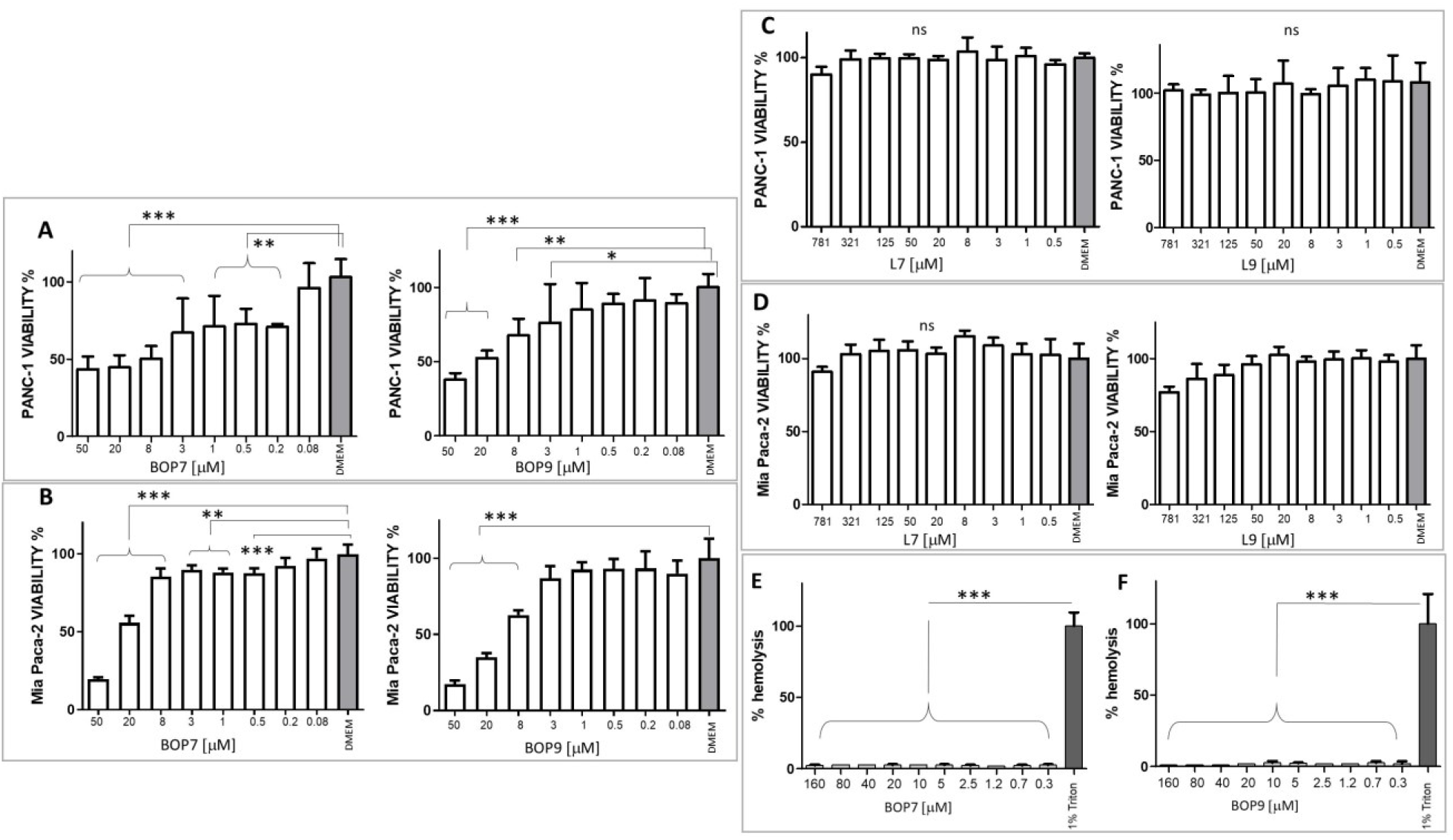
**A**) Viability of PANC-1 cell line after BOP7 treatment. BOP7 IC50: 1.67 × 10^−6^ M, n=6) and BOP9 IC50: 8.3 × 10^−6^ M (n=11); **B**) Viability of Mia PaCa-2 cell line. BOP7 IC50: 2.73 × 10^−5^ M (n=5) and BOP9 IC50: 1.78 × 10^−5^ M (n=5). **C**) Viability of PANC-1 and **D**) Mia PaCa-2 after treatment with L7 and L9; **E**) and **F**) Hemolytic activity of BOP7 and BOP9 (n=3). Data in A-E were analysed with one way ANOVA, Dunnet post test (*** p < 0.0001, ** p<0.01, * p< 0.05). IC50 were calculated in a non-linear fit with log(inhibitor) vs. normalized response variable slope. All graphs are obtained with GraphPad Prism 5.

CHO-K1 cells were chosen as a model of cells of non-cancer origin to test the safety of BOPs. BOP7 showed an IC50 of 5.0 10^−3^ M and BOP9 of 1.6 10^−2^ M, notably less toxic than against pancreatic cell lines. (Supplementary Material 1).

Since many cationic peptides have shown some toxicity towards erythrocytes, BOPs were tested for hemolysis.

The two peptides showed absence of hemolytic activity even at 3 times the highest cytotoxic concentration (Figure 4E-F).

### Effects of BOPs on PANC-1 adhesion and migration

HSPGs play a crucial role in regulating adhesion and migration processes related to invasiveness of cancer cells (28). Through their interaction with HSPGs, BOPs may disrupt adhesion and migration mechanisms in cells. Accordingly, we analysed the effect of the peptides on PANC-1 adhesion to the well. BOP7 and BOP9 at 10 μM inhibited adhesion of PANC-1 when incubated at 37°C for 4 h. BOP7 also produced significant inhibition at 1 μM (Figure 5A). Interference with cell migration was studied in a gap-closure model. Migration of PANC-1 cells was significantly inhibited by BOP7 and BOP9 at 10 μM, especially evident in the case of BOP7 (Figure 5B).

**Figure 5.**
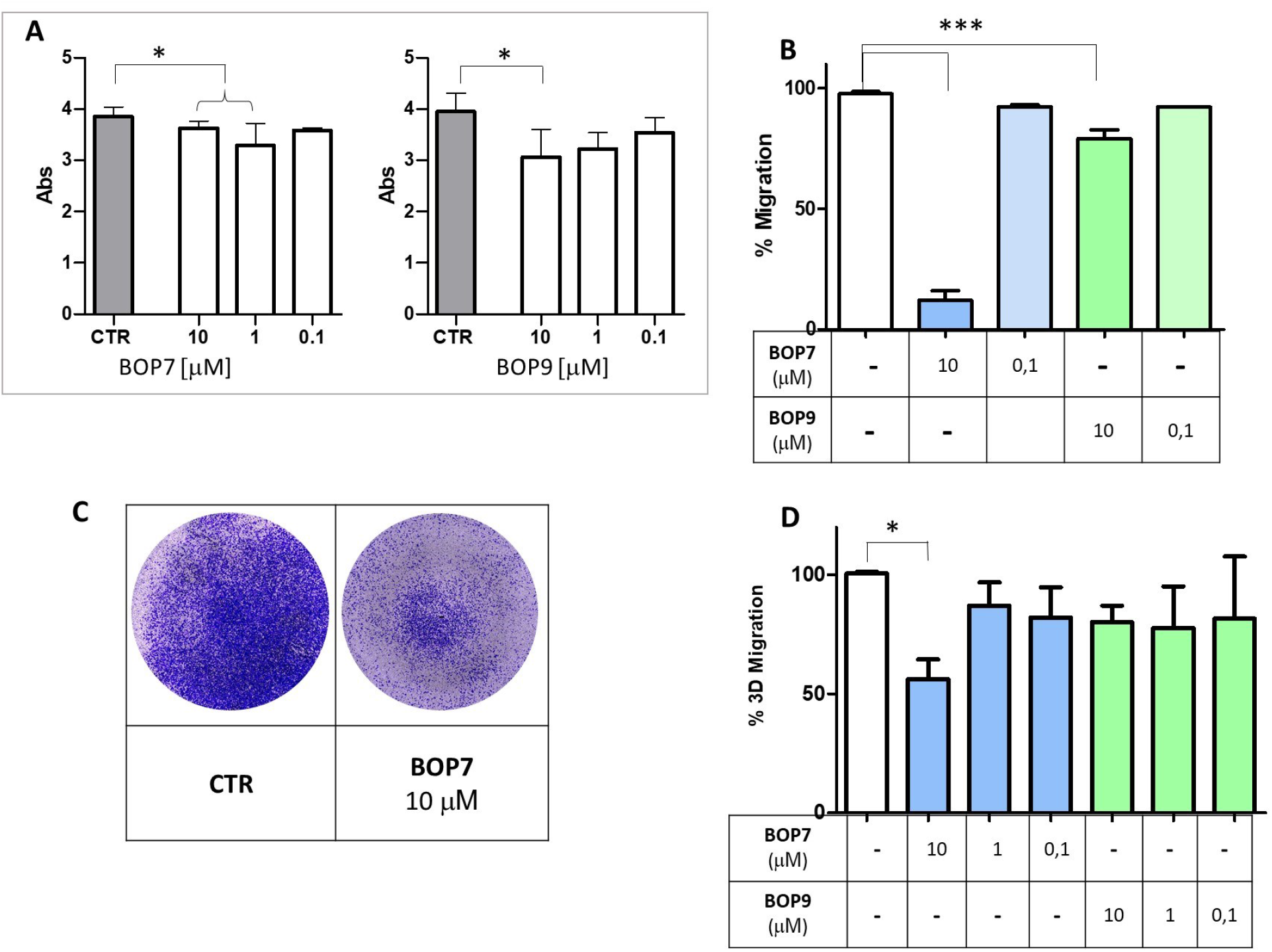
**A**) Inhibition of cell adhesion by BOPs at concentrations 10 μM, 1 μM and 0.1 μM (t test, n=3, *p=0.0243); **B**) Inhibition of cell gap-closure by BOPs at concentrations of 10 μM and 0.1 μM (one way ANOVA, n=3, ***p<0.0001), after 16 h. **C**) Example of 3D migration. **D**) Inhibition of 3D-migration of PANC-1 by BOPs at 10 μM, 1 μM and 0.1 μM (one way ANOVA, n=3, *p=0.0285). All graphs are obtained with GraphPad Prism 5.

To assess whether the effect observed in the two-dimensional migration experiments was correlated with invasiveness *in vitro*, we conducted migration assays using transwell invasion chambers (Figure 5C-D). FBS was used to attract cells and induce their migration from the upper to the lower chamber, passing through the pores of a membrane. In this setting, cells are forced to move, degrade a matrix, collagen type I in this case, and must change shape to squeeze through the pores of the artificial membrane. PANC-1 were seeded on the collagen layer in the upper chamber, and their ability to cross the membrane and reach the lower chamber was measured. BOP7 inhibited PANC-1 3D-migration at a concentration of 10 μM; BOP9 activity was not significant in this setting.

### Immunogenic cell death experiments

To evaluate the ability of BOPs to trigger release of DAMPs, PANC-1 cells were treated with different concentrations of BOPs. The effect was compared to that of two chemotherapeutics used in clinical practice: daunorubicin and irinotecan, both widely considered to be powerful inducers of immunogenic cell death.

### HMGB1 release

Immunogenic cell death inducers trigger release of HMGB1, which by binding to TLR4, promotes the processing and presentation of tumor antigens by inhibiting their premature lysosomal degradation (29-30). HMGB1 release was assessed by ELISA using the supernatants of PANC-1 cells treated for 24 hours with the BOPs and chemotherapeutics at different concentrations (from 300 μM to 2 μM). BOP7 and BOP9 induced greater release of HMGB1 than irinotecan at concentrations of 300 μM and 50 μM, but less than daunorubicin, which proved to be the most effective inducer even at the lowest concentrations (Figure 6A).

**Figure 6.**
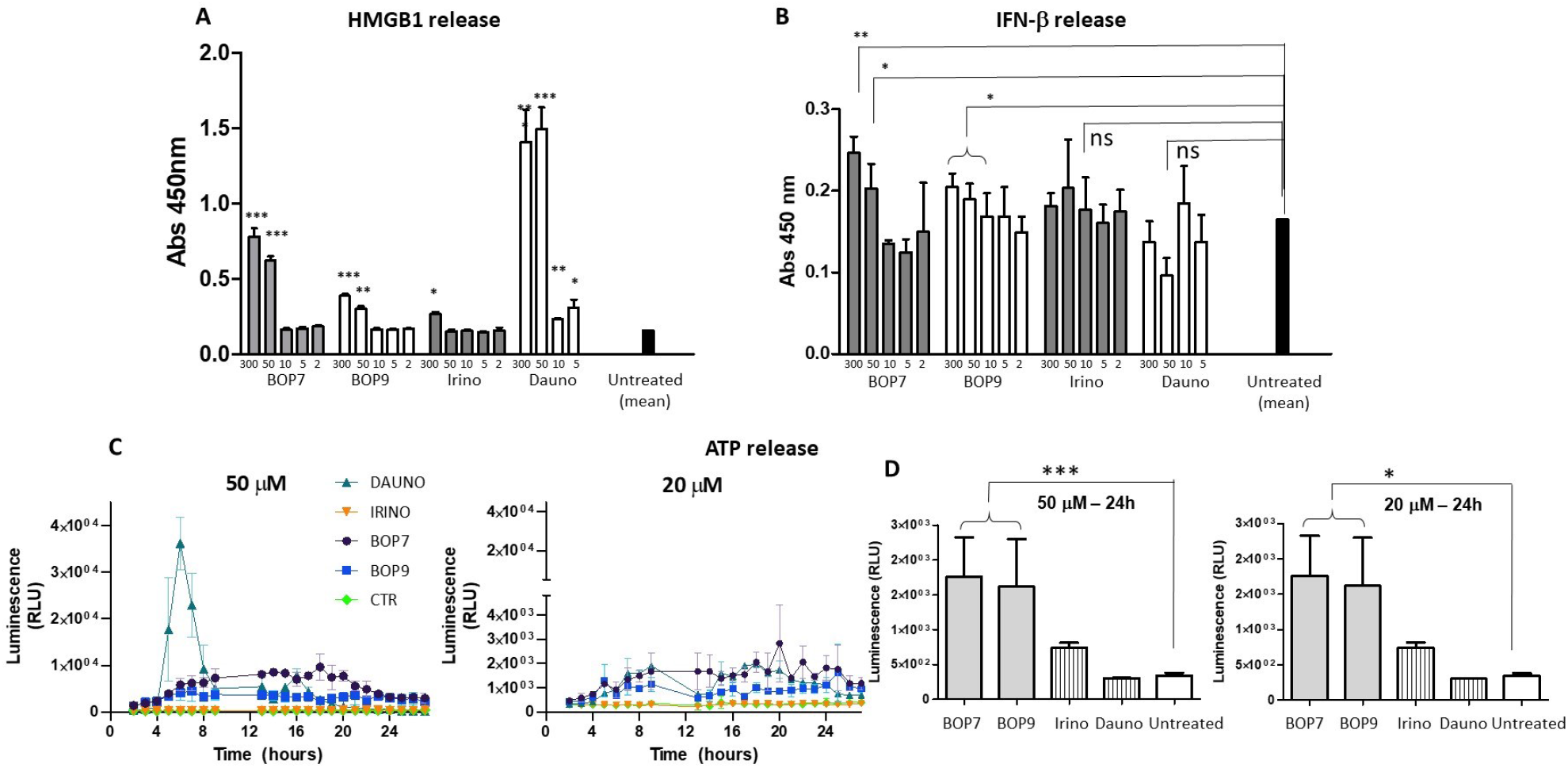
**A**) HMGB1 release by PANC-1 after 24h of treatment with BOPs, daunorubicin (dauno) and irinotecan (irino) (t-tests, unpaired, parametric, two-tailed p-values, every concentration was compared to the mean of untreated cells, *** p < 0.0001, ** p < 0.001, * p < 0.01, n = 3; **B)** IFN-β secretion by PANC-1 after 24h of treatment with BOPs, daunorubicin and irinotecan (t-tests, unpaired, parametric, two-tailed p-values, * p < 0.01, n = 3; **C-D)** ATP release by PANC-1. Bioluminescence at 50 μM and 20 μM BOPs, daunorubicin and irinotecan were plotted against time.

### IFN-β secretion

Immunogenic cell death inducers can stimulate the production of IFN-β type-1 interferon, which acts on its receptor (IFNAR1) and stimulates transcription of specific target genes activating a signal transduction cascade that in turn stimulates T cell recruitment (31-32). IFN-β secretion was assessed by ELISA using the supernatants of PANC-1 cells treated for 24 hours with the different BOPs (Figure 5B). BOP7 and BOP9 at concentrations of 300 μM and 50 μM significantly stimulated IFN-β secretion compared to untreated cells. In contrast, daunorubicin and irinotecan did not result in a statistically significant release of IFN-β. (Figure 6B).

### ATP release

The release of ATP is another hallmark of immunogenic cell death (33). ATP release by PANC-1 cells after treatment with BOPs and chemotherapeutics at different concentrations (from 50 μM to 2 μM), was measured using a bioluminescence test with luciferase. At a concentration of 50 μM, daunorubicin was shown to be the most effective agent at inducing ATP release (Figure 6C). However, the peak of ATP release was reached at 3 hours, and after 6 hours, release returned to baseline. In contrast, at the same concentration, BOP7 and BOP9 stimulated ATP release for longer. After 24 hours, BOP7 and BOP9 at 50 μM and 20 μM released about twice as much ATP as daunorubicin and irinotecan (Figure 6D).

### Mouse model of pancreatic cancer metastases

Considering the tumor-specific anticancer activity of the BOPs, their inhibitory activity in invasiveness/metastatic models *in vitro* and their resistance to circulating proteases, we studied their antitumor effect in a mouse model of metastases.

BALB-c nu/nu mice were injected in the caudal vein with 0.5-1 × 10^6^ PANC1-*luc2* cells (34). PANC1-*luc2* were obtained by transfection of PANC-1 with a plasmid DNA (PGL45LUC2CMVNEO) and proved to have the same susceptibility to BOPs as wild type PANC-1. BOP9 was initially selected for the *in vivo* test because of its slightly lower cytotoxicity against non-cancer cells, compared to BOP7 (IC50 1,6 × 10^−2^ M versus 5,0 × 10^−3^ M, against CHO-K1). Treatments were administered every day for the first 5 days, starting 4 hours after transplant, followed by two days of interval, then three treatment cycles, with two day intervals were repeated for a total of 18 treatments over 32 days (Figure 7A). Tumor growth was monitored using an imaging system (IVIS, PerkinElmer) to capture bioluminescence images of the two groups of mice every 5-6 days (Figure 7B). The images were acquired on anesthetized mice, which were injected with luciferin 15 minutes prior to imaging. BOP9 reduced tumor metastasis growth by 20% with respect to untreated animals (Figure 7C).

**Figure 7.**
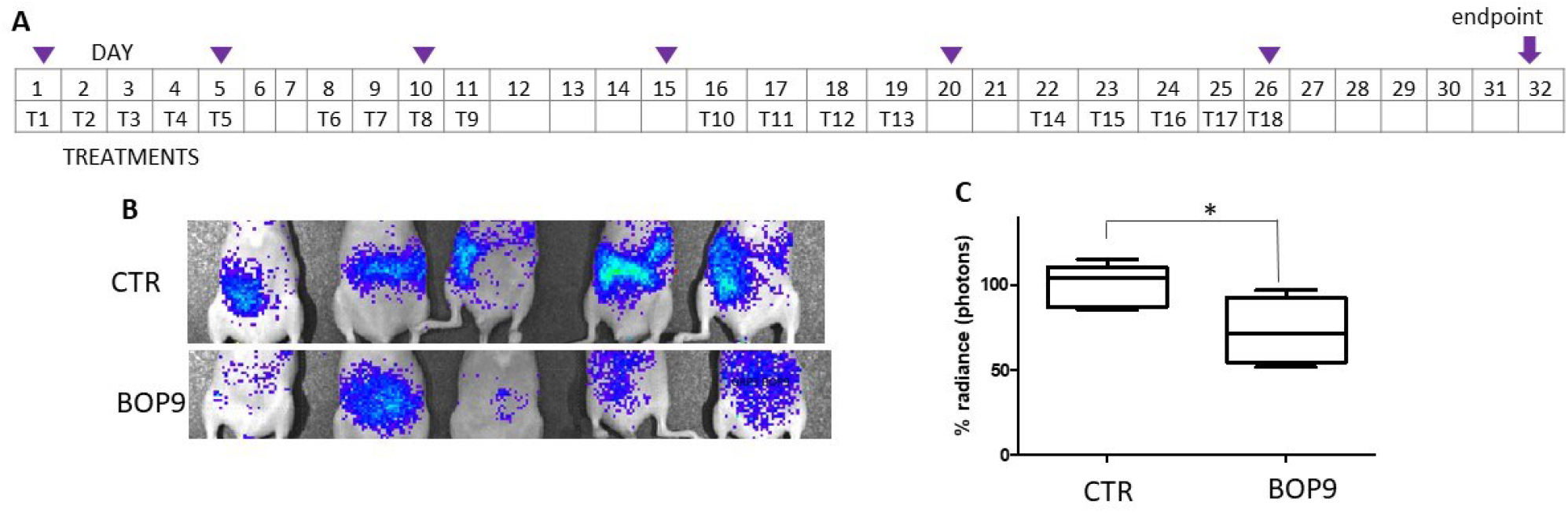
Animal model of metastasis with PANC-1*-luc2*. **A**) Treatment schedule, Tn indicate days of treatment, purple triangles indicate imaging; **B**) representative images of differences in abundance of peritoneal metastases measured as bioluminescence (radiance); **C**) images of the control group (upper panel) and treated group (lower panel) at day 28 (*one tailed unpaired t test, p=0.0200, n=5).

Besides obtaining a reduction in tumor growth, BOPs proved to distribute systemically after intraperitoneal administration and to reach the site of the tumor or metastases.

## DISCUSSION

Tumors with an immunosuppressor microenvironment have poor prognosis. Pancreatic ductal adenocarcinoma is characterized by a prevalence of immunosuppressive infiltrating cells, giving rise to one of the highest mortality rates among all cancer types: the 10-year overall survival rate is approximately 1% (35). Strategies to neutralize the immunosuppressive stroma with high specificity are essential to improve the therapeutic statistics.

Immunogenic cell death (ICD) emerges as a promising solution and several clinical trials have reported a better overall response rate than to chemotherapy alone (36-39). Most studies investigate the ICD-inducing effect of co-treatment with multiple common chemotherapeutics, such as anthracyclines, cyclophosphamide, oxaliplatin, paclitaxel, 5-fluorouracil and immune-check-point inhibitors (40) or passive tumor-targeting immunotherapies, such as trastuzumab (anti-human epidermal growth factor receptor 2).

Peptides hold significant promise in this capacity and are poised to be extensively used in this context in the coming years, owing to their ease of preparation and availability. Anticancer peptides have already been demonstrated to induce ICD (14, 41).

Two peptides, BOP7 and BOP9, showed resistance to proteolysis, unlike their linear analogues. The promising cancer selectivity of the two BOP peptides is attributed to their repeated cationic sequences, which enable multivalent binding to HSPGs bearing multiple anionic sulfation patterns. Cancer cells typically show increased surface negativity due to increased presence of anionic components, such as the sulfate groups and carboxylic moieties of the glycosaminoglycan portion of HSPGs and sialic acid (42). HSPGs are a family of around twenty members, universally expressed in all eukaryotic cells. Many alterations in their expression are reported across different cancer types (43-44). These alterations primarily affect the structure of the glycosaminoglycan chains and the expression levels of the proteoglycans. Both PANC-1 and Mia PaCa-2 cell lines express HSPGs (43-45). The selectivity of the peptides towards cancer cells, observed when comparing PANC-1 or Mia PaCa-2 to the non-cancerous RAW264.7 and CHO cells, is most likely related to an increased expression of HSPGs in the tumor cells, modifications in the GAG chains, or modified arrangement of sulfations on the GAG chains in the pancreatic cancer cell lines (46-47).

This interaction of BOPs with HSPGs not only fosters an anti-metastatic effect in vitro, as shown by reduced adhesion and migration of PANC-1 cancer cells, but also demonstrates promising specific tumor cytotoxicity with low hemolytic activity. Remarkably, the cytotoxicity induced by BOPs triggers release of damage-associated molecular patterns (DAMPs), particularly HMGB1, IFN-β and ATP, by dying cells, and they persist for longer than conventional chemotherapeutic agents such as irinotecan and daunorubicin. The fact that they have a longer-lasting effect on DAMP-release makes

BOPs excellent candidates for combined therapies that leverage immunogenic cell death. The *in vivo* assay in nude mice showed an encouraging ability of BOP9 to inhibit tumor grafting and metastasis growth in a pancreatic cancer model. In addition, BOP9 demonstrated systemic distribution after intraperitoneal administration and thus the ability to effectively reach tumor or metastatic sites, which is a crucial feature for further development.

In light of these promising preliminary results, further studies are needed to precisely define the potential activated pathways of these branched peptides and their mechanism for triggering the release of DAMPs to induce ICD. Additionally, investigating the efficacy of a combined treatment involving BOPs and other clinical standard chemotherapeutics, particularly immune checkpoint inhibitors (6-7), will pave the way for developing new treatment options for non-responding, immunologically silent types of cancers.

## ACKNOWLEDGMENTS

The authors thank Silvia Scali and Stefano Bindi for their contribution. Figures contain elements from Biorender (https://biorender.com), allowed access on the 15/03/2024.

## ADDITIONAL INFORMATION

The authors confirm that the data supporting the findings of this study are available within the article and/or its supplementary materials.

## REFERENCES

1. Di Federico A, Mosca M, Pagani R, Carloni R, Frega G, De Giglio A, Rizzo A, Ricci D, Tavolari S, Di Marco M, Palloni A, Brandi G. Immunotherapy in Pancreatic Cancer: Why Do We Keep Failing? A Focus on Tumor Immune Microenvironment, Predictive Biomarkers and Treatment Outcomes. Cancers (Basel). 2022 May 14;14(10):2429. doi: 10.3390/cancers14102429. PMID: 35626033; PMCID: PMC9139656.

2. Herting CJ, Karpovsky I, Lesinski GB. The tumor microenvironment in pancreatic ductal adenocarcinoma: current perspectives and future directions. Cancer Metastasis Rev. 2021 Sep;40(3):675–689. doi: 10.1007/s10555-021-09988-w. PMID: 34591240.

3. Yin C, Alqahtani A, Noel MS. The Next Frontier in Pancreatic Cancer: Targeting the Tumor Immune Milieu and Molecular Pathways. Cancers (Basel). 2022 May 25;14(11):2619. doi: 10.3390/cancers14112619. PMID: 35681599; PMCID: PMC9179513.

4. Alexandrov LB, Nik-Zainal S, Wedge DC, Aparicio SA, Behjati S, Biankin AV, Bignell GR, Bolli N, Borg A, Børresen-Dale AL, et al. Signatures of mutational processes in human cancer. Nature. 2013 Aug 22;500(7463):415–21. doi: 10.1038/nature12477. Epub 2013 Aug Erratum in: Nature. 2013 Oct 10;502(7470):258. Imielinsk, Marcin [corrected to Imielinski, Marcin]. PMID: 23945592; PMCID: PMC3776390.

5. Hou W, Yang B, Zhu H. Nanoparticle-Based Therapeutic Strategies for Enhanced Pancreatic Ductal Adenocarcinoma Immunotherapy. Pharmaceutics. 2022 Sep 24;14(10):2033. doi: 10.3390/pharmaceutics14102033. PMID: 36297467; PMCID: PMC9607590.

6. Schizas D, Charalampakis N, Kole C, Economopoulou P, Koustas E, Gkotsis E, Ziogas D, Psyrri A, Karamouzis MV. Immunotherapy for pancreatic cancer: A 2020 update. Cancer Treat Rev. 2020 Jun;86:102016. doi: 10.1016/j.ctrv.2020.102016. Epub 2020 Mar 25. PMID: 32247999.

7. Muller M, Haghnejad V, Schaefer M, Gauchotte G, Caron B, Peyrin-Biroulet L, Bronowicki JP, Neuzillet C, Lopez A. The Immune Landscape of Human Pancreatic Ductal Carcinoma: Key Players, Clinical Implications, and Challenges. Cancers (Basel). 2022 Feb 16;14(4):995. doi: 10.3390/cancers14040995. PMID: 35205742; PMCID: PMC8870260.

8. Kroemer G, Galassi C, Zitvogel L, Galluzzi L. Immunogenic cell stress and death. Nat Immunol. 2022 Apr;23(4):487–500. doi: 10.1038/s41590-022-01132-2. Epub 2022 Feb 10. PMID: 35145297.

9. Zhou J, Wang G, Chen Y, Wang H, Hua Y, Cai Z. Immunogenic cell death in cancer therapy: Present and emerging inducers. J Cell Mol Med. 2019 Aug;23(8):4854–4865. doi: 10.1111/jcmm.14356. Epub 2019 Jun 18. PMID: 31210425; PMCID: PMC6653385.

10. Galluzzi L, Buqué A, Kepp O, Zitvogel L, Kroemer G. Immunogenic cell death in cancer and infectious disease. Nat Rev Immunol. 2017 Feb;17(2):97–111. doi: 10.1038/nri.2016.107. Epub 2016 Oct 17. PMID: 27748397.

11. Yamazaki T, Buqué A, Ames TD, Galluzzi L. PT-112 induces immunogenic cell death and synergizes with immune checkpoint blockers in mouse tumor models. Oncoimmunology. 2020 Feb 11;9(1):1721810. doi: 10.1080/2162402X.2020.1721810. PMID: 32117585; PMCID: PMC7028345.

12. Gomes-da-Silva LC, Zhao L, Bezu L, Zhou H, Sauvat A, Liu P, Durand S, Leduc M, Souquere S, Loos F, Mondragón L, Sveinbjørnsson B, Rekdal Ø, Boncompain G, Perez F, Arnaut LG, Kepp O, Kroemer G. Photodynamic therapy with redaporfin targets the endoplasmic reticulum and Golgi apparatus. EMBO J. 2018 Jul 2;37(13):e98354. doi: 10.15252/embj.201798354. Epub 2018 May 28. PMID: 29807932; PMCID: PMC6028029.

13. Shekarian T, Sivado E, Jallas AC, Depil S, Kielbassa J, Janoueix-Lerosey I, Hutter G, Goutagny N, Bergeron C, Viari A, Valsesia-Wittmann S, Caux C, Marabelle A. Repurposing rotavirus vaccines for intratumoral immunotherapy can overcome resistance to immune checkpoint blockade. Sci Transl Med. 2019 Oct 23;11(515):eaat5025. doi: 10.1126/scitranslmed.aat5025. PMID: 31645452.

14. Ghaly G, Tallima H, Dabbish E, Badr ElDin N, Abd El-Rahman MK, Ibrahim MAA, Shoeib T. Anti-Cancer Peptides: Status and Future Prospects. Molecules. 2023 Jan 23;28(3):1148. doi: 10.3390/molecules28031148. PMID: 36770815; PMCID: PMC9920184.

15. Tornesello AL, Borrelli A, Buonaguro L, Buonaguro FM, Tornesello ML. Antimicrobial Peptides as Anticancer Agents: Functional Properties and Biological Activities. Molecules. 2020 Jun 19;25(12):2850. doi: 10.3390/molecules25122850. PMID: 32575664; PMCID: PMC7356147.

16. Liu H, Shen W, Liu W, Yang Z, Yin D, Xiao C. From oncolytic peptides to oncolytic polymers: A new paradigm for oncotherapy. Bioact Mater. 2023 Aug 14;31:206–230. doi: 10.1016/j.bioactmat.2023.08.007. PMID: 37637082; PMCID: PMC10450358.

17. Galluzzi L, Vitale I, Warren S, Adjemian S, Agostinis P, Martinez AB, Chan TA, Coukos G, Demaria S, Deutsch E, et al. Consensus guidelines for the definition, detection and interpretation of immunogenic cell death. J Immunother Cancer. 2020 Mar;8(1):e000337. doi: 10.1136/jitc-2019-000337. Erratum in: J Immunother Cancer. 2020 May;8(1): PMID: 32209603; PMCID: PMC7064135.

18. Monroc S, Badosa E, Besalú E, Planas M, Bardají E, Montesinos E, Feliu L. Improvement of cyclic decapeptides against plant pathogenic bacteria using a combinatorial chemistry approach. Peptides. 2006 Nov;27(11):2575–84. doi: 10.1016/j.peptides.2006.05.001. Epub 2006 Jun 9. PMID: 16762457

19. Feliu L, Oliveras G, Cirac AD, Besalú E, Rosés C, Colomer R, Bardají E, Planas M, Puig T. Antimicrobial cyclic decapeptides with anticancer activity. Peptides. 2010 Nov;31(11):2017–26. doi: 10.1016/j.peptides.2010.07.027. Epub 2010 Aug 11. PMID: 20708052

20. Chen CJ, Tsai KC, Kuo PH, Chang PL, Wang WC, Chuang YJ, Chang MD. A Heparan Sulfate-Binding Cell Penetrating Peptide for Tumor Targeting and Migration Inhibition. Biomed Res Int. 2015;2015:237969. doi: 10.1155/2015/237969. Epub 2015 May 3. PMID: 26064887; PMCID: PMC4433633

21. Brunetti J, Falciani C, Lelli B, Minervini A, Ravenni N, Depau L, Siena G, Tenori E, Menichetti S, Pini A, et al. Neurotensin branched peptide as a tumor-targeting agent for human bladder cancer. Biomed Res Int. 2015;2015:173507. doi: 10.1155/2015/173507. Epub 2015 Apr 23. PMID: 25984525; PMCID: PMC4423026.

22. Falciani C, Brunetti J, Lelli B, Ravenni N, Lozzi L, Depau L, Scali S, Bernini A, Pini A, Bracci L. Cancer selectivity of tetrabranched neurotensin peptides is generated by simultaneous binding to sulfated glycosaminoglycans and protein receptors. J Med Chem. 2013 Jun 27;56(12):5009–18. doi: 10.1021/jm400329p. Epub 2013 Jun 6. PMID: 23713525.

23. Li X, Zhong H, Zheng S, Mu J, Yu N, Guo S. Tumor-penetrating iRGD facilitates penetration of poly(floxuridine-ketal)-based nanomedicine for enhanced pancreatic cancer therapy. J Control Release. 2024 May;369:444–457. doi: 10.1016/j.jconrel.2024.04.004. Epub 2024 Apr 6. PMID: 38575076

24. Nielsen M, Monberg T, Sundvold V, Albieri B, Hovgaard D, Petersen MM, Krarup-Hansen A, Met Ö, Camilio K, Clancy T, Stratford R, Sveinbjornsson B, Rekdal Ø, Junker N, Svane IM. LTX-315 and adoptive cell therapy using tumor-infiltrating lymphocytes generate tumor specific T cells in patients with metastatic soft tissue sarcoma. Oncoimmunology. 2023 Dec 7;13(1):2290900. doi: 10.1080/2162402X.2023.2290900. PMID: 38125722; PMCID: PMC10732595

25. Bracci L, Falciani C, Lelli B, Lozzi L, Runci Y, Pini A, De Montis MG, Tagliamonte A, Neri P. Synthetic peptides in the form of dendrimers become resistant to protease activity. J Biol Chem. 2003 Nov 21;278(47):46590–5. doi: 10.1074/jbc.M308615200. Epub 2003 Sep 12. PMID: 12972419.

26. Falciani C, Lozzi L, Pini A, Corti F, Fabbrini M, Bernini A, Lelli B, Niccolai N, Bracci L. Molecular basis of branched peptides resistance to enzyme proteolysis. Chem Biol Drug Des. 2007 Mar;69(3):216–21. doi: 10.1111/j.1747-0285.2007.00487.x. PMID: 17441908.

27. Paul S, Verma S, Chen YC. Peptide Dendrimer-Based Antibacterial Agents: Synthesis and Applications. ACS Infect Dis. 2024 Mar 1. doi: 10.1021/acsinfecdis.3c00624. Epub ahead of print. PMID: 38428037.

28. Esko JD, Rostand KS, Weinke JL. Tumor formation dependent on proteoglycan biosynthesis. Science. 1988 Aug 26;241(4869):1092–6. doi: 10.1126/science.3137658. PMID: 3137658.

29. Yang H, Wang L. Heparan sulfate proteoglycans in cancer: Pathogenesis and therapeutic potential. Adv Cancer Res. 2023;157:251–291. doi: 10.1016/bs.acr.2022.08.001. Epub 2022 Aug 29. PMID: 36725112.

30. Apetoh L, Ghiringhelli F, Tesniere A, Obeid M, Ortiz C, Criollo A, Mignot G, Maiuri MC, Ullrich E, Saulnier P, et al. Toll-like receptor 4-dependent contribution of the immune system to anticancer chemotherapy and radiotherapy. Nat Med. 2007 Sep;13(9):1050–9. doi: 10.1038/nm1622. Epub 2007 Aug 19. PMID: 17704786.

31. Apetoh L, Ghiringhelli F, Tesniere A, Criollo A, Ortiz C, Lidereau R, Mariette C, Chaput N, Mira JP, Delaloge S, et al. The interaction between HMGB1 and TLR4 dictates the outcome of anticancer chemotherapy and radiotherapy. Immunol Rev. 2007 Dec;220:47–59. doi: 10.1111/j.1600-065X.2007.00573.x. PMID: 17979839.

32. Sistigu A, Yamazaki T, Vacchelli E, Chaba K, Enot DP, Adam J, Vitale I, Goubar A, Baracco EE, Remédios C, et al. Cancer cell-autonomous contribution of type I interferon signaling to the efficacy of chemotherapy. Nat Med. 2014 Nov;20(11):1301–9. doi: 10.1038/nm.3708. Epub 2014 Oct 26. PMID: 25344738.;

33. Ahmed A, Tait SWG. Targeting immunogenic cell death in cancer. Mol Oncol. 2020 Dec;14(12):2994–3006. doi: 10.1002/1878-0261.12851. Epub 2020 Dec 1. PMID: 33179413; PMCID: PMC7718954.

34. Krysko O, Løve Aaes T, Bachert C, Vandenabeele P, Krysko DV. Many faces of DAMPs in cancer therapy. Cell Death Dis. 2013 May 16;4(5):e631. doi: 10.1038/cddis.2013.156. PMID: 23681226; PMCID: PMC3674363.

35. Yuan P, He XH, Rong YF, Cao J, Li Y, Hu YP, Liu Y, Li D, Lou W, Liu MF. KRAS/NF-κB/YY1/miR-489 Signaling Axis Controls Pancreatic Cancer Metastasis. Cancer Res. 2017 Jan 1;77(1):100–111. doi: 10.1158/0008-5472.CAN-16-1898. Epub 2016 Oct 28. Erratum in: Cancer Res. 2020 Sep 15;80(18):4022. PMID: 27793842

36. Siegel RL, Miller KD, Fuchs HE, Jemal A. Cancer statistics, 2022. CA Cancer J Clin. 2022 Jan;72(1):7–33. doi: 10.3322/caac.21708. Epub 2022 Jan 12. PMID: 35020204.

37. Sprooten J, Laureano RS, Vanmeerbeek I, Govaerts J, Naulaerts S, Borras DM, Kinget L, Fucíková J, Špíšek R, Jelínková LP, et al. Trial watch: chemotherapy-induced immunogenic cell death in oncology. Oncoimmunology. 2023 Jun 3;12(1):2219591. doi: 10.1080/2162402X.2023.2219591. eCollection 2023.

38. Zhang B, Li N, Gao J, Zhao Y, Jiang J, Xie S, Zhang C, Zhang Q, Liu L, Wang Z, Ji D, Wu L, Ren R. Targeting of focal adhesion kinase enhances the immunogenic cell death of PEGylated liposome doxorubicin to optimize therapeutic responses of immune checkpoint blockade. J Exp Clin Cancer Res. 2024 Feb 19;43(1):51. doi: 10.1186/s13046-024-02974-4. PMID: 38373953; PMCID: PMC10875809;

39. Islam MR, Patel J, Back PI, Shmeeda H, Kallem RR, Shudde C, Markiewski M, Putnam WC, Gabizon AA, La-Beck NM. Pegylated Liposomal Alendronate Biodistribution, Immune Modulation, and Tumor Growth Inhibition in a Murine Melanoma Model. Biomolecules. 2023 Aug 26;13(9):1309. doi: 10.3390/biom13091309. PMID: 37759709; PMCID: PMC10527549;

40. Galassi C, Klapp V, Yamazaki T, Galluzzi L. Molecular determinants of immunogenic cell death elicited by radiation therapy. Immunol Rev. 2024 Jan;321(1):20–32. doi: 10.1111/imr.13271. Epub 2023 Sep 7. PMID: 37679959.

41. Jiang Y-Z, Liu Y, Xiao Y, Hu X, Jiang L, Zuo W-J, Ma D, Ding J, Zhu X, Zou J, et al. Molecular subtypi ng and genomic profiling expand precision medicine in refractory metastatic triple-negative breast cancer: the FUTURE trial. Cell Res. 2021;31:178–186. doi:10.1038/s41422-020-0375-9

42. Pasquereau-Kotula E, Habault J, Kroemer G, Poyet J-L, Ma W-L. The anticancer peptide RT53 induces immunogeniccell death. PLos One. 2018;13:e0201220. doi:10.1371/journal.pone.0201220

43. Liu H, Shen W, Liu W, Yang Z, Yin D, Xiao C. From oncolytic peptides to oncolytic polymers: A new paradigm for oncotherapy. Bioact Mater. 2023 Aug 14;31:206–230. doi: 10.1016/j.bioactmat.2023.08.007. PMID: 37637082; PMCID: PMC10450358.

44. Furini S, Falciani C. Expression and Role of Heparan Sulfated Proteoglycans in Pancreatic Cancer. Front Oncol. 2021 Jun 25;11:695858. doi: 10.3389/fonc.2021.695858. PMID: 34249755; PMCID: PMC8267412 https://www.proteinatlas.org/search/proteoglycan accessed on the17 July 2024

45. Depau L, Brunetti J, Falciani C, Mandarini E, Riolo G, Zanchi M, Karousou E, Passi A, Pini A, Bracci L. Heparan Sulfate Proteoglycans Can Promote Opposite Effects on Adhesion and Directional Migration of Different Cancer Cells. J Med Chem. 2020 Dec 24;63(24):15997–16011. doi: 10.1021/acs.jmedchem.0c01848. Epub 2020 Dec 7. PMID: 33284606

46. Kleeff J, Ishiwata T, Kumbasar A, Friess H, Büchler MW, Lander AD, et al. The Cell-Surface Heparan Sulfate Proteoglycan Glypican-1 Regulates Growth Factor Action in Pancreatic Carcinoma Cells and Is Overexpressed in Human Pancreatic Cancer. J Clin Invest (1998) 102(9):1662–73. doi: 10.1172/JCI4105

47. Uhlen M, Zhang C, Lee S, Sjöstedt E, Fagerberg L, Bidkhori G, et al. A Pathology Atlas of the Human Cancer Transcriptome. Science (2017) 357 (6352):eaan2507. doi: 10.1126/science.aan2507

