## Supplementary Figure 1 for "Branched Oncolytic Peptides Target HSPGs, Inhibit Metastasis, and Trigger the Release of Molecular Determinants of Immunogenic Cell Death in Pancreatic Cancer"

SUPPLEMENTARY MATERIAL

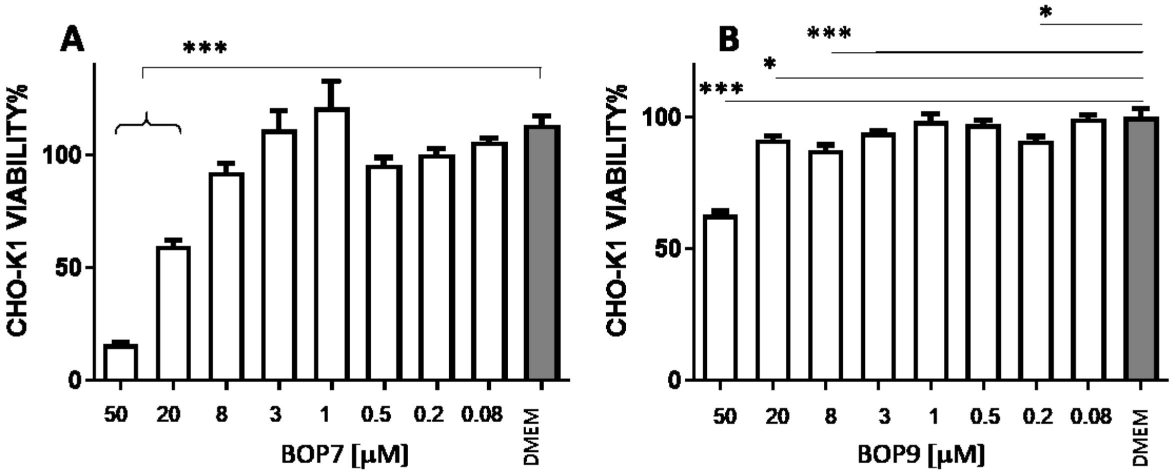


**Supplementary Material 1**. Viability of CHO-K1 cell line after treatment with BOP7 (A) IC50 5.0 x $10^{-3}$ M (n=5) IC50 and BOP9(B) IC50: 1.6 x $10^{-2}$ M (n=5). Data were analysed with one way ANOVA, Dunnet post test (*** $p < 0.0001$, * $p< 0.05$). IC50 were calculated in a non-linear fit variable slope. All graphs are obtained with GraphPad Prism 5.
